# Inbreeding reduces fitness of seed beetles under thermal stress

**DOI:** 10.1101/2021.05.05.442709

**Authors:** Edward Ivimey-Cook, Sophie Bricout, Victoria Candela, Alexei A. Maklakov, Elena C. Berg

## Abstract

Human-induced environmental change can influence populations both at the global level through climatic warming and at the local level through habitat fragmentation. As populations become more isolated, they can suffer from high levels of inbreeding which contributes to a reduction in fitness, termed inbreeding depression. However, it is still unclear if this increase in homozygosity also results in a corresponding increase in sensitivity to stressful conditions, which could intensify the already detrimental effects of environmental warming. Here, in a fully factorial design, we assessed the life-long impact of increased mutation load and elevated temperature on key life history traits in the seed beetle, *Callosobruchus maculatus*. We found that beetles raised at higher temperatures had far reduced fitness and survival than beetles from control temperatures. Importantly, these negative effects were exacerbated in inbred beetles as a result of increased mutation load, with further detrimental effects manifesting on individual hatching probability and lifetime reproductive success. These results reveal the harmful impact that increasing temperature and likelihood of habitat fragmentation due to anthropogenetic changes in environmental conditions could have on populations of organisms worldwide.

## Introduction

The Earth’s average annual temperature has risen by approximately 0.85°C over the past 100 years (Pereira *et al.*, 2012; Pachauri *et al.*, 2014) with the current rate of warming nearly double that of previous decades (Rosenzweig *et al.*, 2008; Pereira *et al.*, 2012; Pachauri *et al.*, 2014). One of the major contributors to this rise in annual temperature is anthropogenic greenhouse gas emissions, which have caused more than half of the observed increase in global average surface temperature from 1951 to 2010 (Pereira *et al.*, 2012; Pachauri *et al.*, 2014). This unprecedented rise in temperature is already affecting natural systems (Pereira *et al.*, 2012; Pachauri *et al.*, 2014; Trisos *et al.*, 2020), driving many organisms to either adapt, move, or go extinct (Holt, 1990; Pereira *et al.*, 2012; Trisos *et al.*, 2020).

In particular, a warmer and more unpredictable climate has forced many organisms, from both terrestrial and marine environments, to shift geographic ranges, alter seasonal activities or migration patterns, or change interactions with other species (Barnett *et al.*, 2001; Root *et al.*, 2003). For instance, it is predicted that many terrestrial and freshwater species will significantly alter range boundaries and move polewards in response to anthropogenic warming as thermal tolerances are likely to be exceeded nearer the equator (Hickling *et al.*, 2006; Thomas, 2010). This shift in geographical range may also lead to corresponding changes in species interactions within ecosystems. For instance, a review comprising data from 688 published studies found significant, multitrophic effects of global environmental change acting on both mutualistic and antagonist interactions among species within an ecosystem (Tylianakis *et al.*, 2008). Species interactions could also change as a result of altered migration patterns. For example, in response to warmer winters, several bird species have substantially reduced the migration distance between breeding and overwintering grounds (Visser *et al.*, 2009).

A wealth of literature has also revealed how changes in climatic conditions can have cascading effects on the life history and ability of an organism to adapt to shifts in phenology (Davis & Shaw, 2001; Gottfried *et al.*, 2012; Norberg *et al.*, 2012; Pearson *et al.*, 2014; Muñoz *et al.*, 2015; Seebacher *et al.*, 2015). Some species are able to adapt sufficiently by undergoing rapid evolutionary change. For example, in response to a five-year period of drought, the southern Californian plant species *Brassica rapa* shifted to an earlier flowering time and increased the overall duration of flowering. Subsequently, this change in flowering time then led to an increase in individual fitness as a result of escaping the harsh conditions of late-season drought (Franks & Weis, 2008). Other species, which have been unable to adapt as quickly, have seen substantial population declines. For example, in several European bird species, warmer temperatures have resulted in phenological mismatch between breeding opportunities and food peaks (Visser *et al.*, 1998, 2012; Both *et al.*, 2006; Jiguet *et al.*, 2007).

The ability of an organism to undergo rapid adaptation to novel ecological conditions such as elevated temperature is reliant on the existence of standing genetic variation within a population (Davis & Shaw, 2001; Orr & Betancourt, 2001; Blows & Hoffmann, 2005; Willi *et al.*, 2006; Berger *et al.*, 2020). Therefore, a reduction in genetic diversity could restrict the evolvability of populations to environmental stochasticity. Climate warming and increased anthropogenic land use change (Opdam & Wascher, 2004; Liao & Reed, 2009) have led to habitat fragmentation (and habitat loss), which can induce genetic constraints on adaptation by increasing the levels of inbreeding (Leimu *et al.*, 2006) as populations become more isolated. This increase in genetic homozygosity within a population often results in a significant reduction to survival and fertility through the expression of deleterious, recessive mutations (Keller & Waller, 2002; Charlesworth & Willis, 2009), termed inbreeding depression (Charlesworth & Charlesworth, 1987).

In the wild, inbreeding depression is both widespread and variable in magnitude within and between populations (Keller & Waller, 2002; Huisman *et al.*, 2016). Importantly for conservation biologists this increase in mutation load (Kirkpatrick & Jarne, 2000) and loss of genetic diversity (Gibbs, 2001) could potentially exaggerate a population’s sensitivity to environmental stress and increase the likelihood of extinction (Bijlsma *et al.*, 1999; Fox *et al.*, 2006, 2011; Franke & Fischer, 2015).

Inbred individuals may have a heightened sensitivity to increased environmental stress, through factors such as temperature, competition, nutrition, exposure to harmful chemicals, parasitism and desiccation. This sensitivity has been investigated in several species to date (See Armbruster & Reed, 2005, Agrawal & Whitlock, 2010 and Fox & Reed, 2011), including models systems such as the seed beetle *Callosobruchus maculatus* (Fox *et al.*, 2006, 2011; Fox & Stillwell, 2009; Fox & Reed, 2010) and the fruit fly *Drosophila melanogaster* (Yun & Agrawal, 2014). However, crucially for conservation research, the link between thermal stress and inbreeding depression remains unclear. In addition, a recent study by Yun and Agrawal (2014) highlighted that much of the link between environmental stress and inbreeding depression could be a result of density dependence (competition stress) driving the interaction.

Despite this, a recent study has shown that increasing temperature results in significantly more genome-wide *de novo* mutations (Berger *et al.*, 2020). However, empirical support for a corresponding increase in inbreeding depression owing to the accumulation of these thermal stress-induced mutations is varied. For instance, in a series of studies, Fox *et al.* found that inbreeding depression on larval developmental traits either increased (Fox & Reed, 2011) or decreased (Fox *et al.*, 2011) in environments of high thermal stress. In particular, the latter experiment found that inbred individuals were detrimentally affected at the more benign temperature of 20°C as opposed to the higher, elevated temperatures in the previous experiment (Fox *et al.*, 2011). Not only are the results from these experiments seemingly contradictory but they are also solely focused on measuring inbreeding depression manifesting on larval developmental traits (survival and generation time) under *developmental* stress.

Therefore, to fully understand the interaction between environmental stress and inbreeding depression, it is necessary to study its effect on both survival and fecundity. In addition, exposing individuals to stress across the entirety of their lifespan, and not just the developmental period, would more accurately reflect changes to environment predicted as a result of global climatic change. In light of this and in order to address the paucity of data surrounding inbreeding depression and thermal stress, we examined the impact of inbreeding and mutation load on the lifespan and fitness of the model system, *C. maculatus*, when exposed to two different *lifelong* rearing temperatures, one stressful and one benign.

## Methods

### Study system

The seed beetle (*C. maculatus*), native to Africa and Asia, is an agricultural pest that infests legumes in warehouses and in the field. Females lay their eggs on the surface of host seeds (Messina, 1991; Fox *et al.*, 2006). Eggs hatch 4–5 days later and larvae burrow into the seed (Fox *et al.*, 2006). Larvae develop inside the bean, and the beetles emerge as reproductively mature adults after around 23-27 days. *C. maculatus* beetles are facultatively aphageous – that is, they are able to acquire all the water and food resources they need from the bean during larval development and do not require additional resources as adults (Messina & Slade, 1999). In part because of the ease of laboratory rearing, *C. maculatus* has become a model organism for the study of sex differences in life history evolution (Fox, 1994; Fox *et al.*, 2006, 2007; Bilde *et al.*, 2009; Maklakov & Fricke, 2009; Fritzsche & Arnqvist, 2013).

The study population “South India USA” originated from an outbred stock population that was collected from infested mung beans (*Vigna radiata*) in Tirunelveli, India, in 1979. They were then moved by C. W. Fox to the University of Kentucky, USA, then to Uppsala University in 1992, and finally to the American University of Paris in 2015. The stock population is kept at aphagy (no food or water) in 1L jars with 150g of mung beans, and approximately 250 newly hatched beetles are transferred to new jars with fresh beans every 23-24 days on a continual basis. The beetles are maintained in climate chambers at 29°C, 50% relative humidity and a 12:12 h light:dark cycle. These laboratory conditions closely resemble their natural conditions, since their life history is adapted to a storage environment (Messina, 1991; Fox, 1994).

### Experimental groups

From the base population, we created four experimental treatments that differed in level of inbreeding as well as rearing temperature. The first step was to generate “inbred” (I) beetles, which were the offspring of full sibling pairs. To do this, fertilized beans were transferred from the stock jars to virgin chambers (aerated plastic culture plates with a separate well for each individual) and monitored daily. Approximately 24 hours after hatch, one male and one female were randomly paired together and placed in a 60-mm Petri dish with approximately 80 beans (N = 50). All adults were removed 48 hours later, and larvae were left to develop.

Before the next generation hatched, 48 fertilized beans were moved from each Petri dish to individually labelled 48-well virgin chamber plates, which were monitored daily. Approximately 24 hours after hatch, one sister and one brother from each 48-well plate were placed together into a 60-mm dish with approximately 70 beans (N = 50 inbred pairs). Meanwhile, we created 50 “outbred” (O) pairings between randomly selected one-day-old males and females that had hatched out of fertilized beans (isolated in virgin chambers) from the background population. All of the inbred and outbred pairs were created on the same day.

Next, we created the four different treatment groups: outbred at the “control” temperature of 29°C (OC), outbred at the “elevated” temperature of 36°C (OE), inbred at 29°C (IC), and inbred at 36°C (IE). To do this, approximately 24 hours after pairing the beetles as described above, ten fertilized beans from each petri (N = 50 inbred and 50 outbred dishes) were randomly selected and placed into two carefully labeled virgin chambers, five beans per virgin chamber. We selected only those beans that had eggs on them that appeared to be viable (clear, round and regularly shaped, firmly attached to the bean). One of the plates was placed into a climate chamber kept at the control temperature (29°C), and the other plate was placed in a chamber set to “elevated” temperature (36°C). This higher temperature was selected because it represents the upper limit of what the beetles can withstand without devastating impacts on fertility or lifespan (Rogell *et al.*, 2014). Humidity and light: dark cycles were kept the same for both chambers: 50% humidity and 12:12 h light:dark. Virgin chambers were monitored daily.

### Daily fecundity and lifespan assays

We monitored the virgin chambers every day and recorded the hatch date and sex of all eclosed offspring from the four treatments. One day after hatch, we paired the offspring with a one-day old beetle of the opposite sex from the background population. Similar to previous steps of the experiment, virgin background beetles were generated by putting fertilized beans from the control jars into virgin chambers, and hatch was monitored daily. Pairs were moved at the same time every day from one Petri dish to another for five days.

On the day of pairing (D0), the male and female were placed in a 60-mm Petri with 65 beans. Females can lay up to 65 eggs per day (E. C. Berg, unpublished data), and we wanted to provide enough beans so that no more than one egg would be laid on each bean. On subsequent days (D1, D2, D3, and D4+), pairs were moved to 35-mm Petri dishes with 30-50 beans (egg-laying declines with age). Once the pairs were moved to the final dish in the series, they were monitored daily. If at any point the female was found dead, pairs were obviously not transferred further. All dead individuals were removed immediately, and dates of death were recorded.

To calculate daily fecundity, we recorded the number of eclosed offspring per dish per target individual. Approximately 35 days after eggs were laid, we froze the dishes to facilitate counting of eclosed offspring.

### Statistical analyses

All analyses were performed using R v4.0.3 (R Core Team, 2019). Four distinct measures of reproduction were analysed using the glmmTMB v1.0.2.9000 package (Brooks *et al.*, 2017; Magnusson *et al.*, 2019) and contained the main effects of “Breeding status” (inbred or outbred) and “Temperature regime” (control or elevated) and the subsequent higher-order interaction. In addition, all models contained the random effect of “Parent ID” in order to account for pseudoreplication of individuals from the same parent. For age-specific reproduction, additional fixed effects of Day and Day^2^ and an additional random effect of “Individual ID” was added, nested within “Parent ID”, in order to account for repeatedly measuring the same individual over time.

Whilst the fixed and random effect structure remained similar for each measure, the distributions of the responses differed slightly. 1) Hatching success was a binary response, where individuals either hatched (1) or did not (0). 2) For both age-specific reproduction and lifetime reproductive success (LRS), data was analysed in a two-step process. Firstly, a full Poisson model and a Poisson model with an observation level random effect was fitted and the residuals simulated using the DHARMa v0.3.3.0 package (Hartig, 2020). If zero-inflation was detected within these residuals, an additional zero-inflation component and a variety of error distributions were fitted. Model selection was then performed to select the best fitting error distribution and zero-inflation parameters for each measure, chosen as the model with the lowest Akaike’s information criterion (AIC). 3) The last measure was individual fitness, or λ*ind*, which represented the dominant eigenvalue of an age-structured Leslie matrix (Leslie, 1945) calculated using the popbio v2.7 package (Stubben & Milligan, 2007). For each matrix, the top row denoted age-specific fertility whilst the subdiagonal represented survival probability from age *t* to *t+1*. 18-32 days were also added to the start of the fertility schedule which corresponded to egg-adult development time under the various breeding and temperature treatments. These individual fitness values were then analysed with a similar model structure to above, albeit with a Gaussian error structure.

For each measure, the overall effect of “Breeding status”, “Temperature regime”, and the interaction between the two, was identified using the Anova function from the car v3.0-10 package. In addition, data was either visualised using the ggplot2 v3.3.3 package (Wickham, 2009) or on bootstrapped estimation plots from the dabestR v0.3.0 package (Ho *et al.*, 2019). Estimated marginal means were reported using the emmeans v1.5.5-1 package (Lenth *et al.*, 2019).

Lastly, we analysed how “Temperature regime” and “Breeding status” influenced survival and lifespan. For this we used Mixed Effects Cox Proportional Hazards Models from the coxme package v2.2-16 (Therneau, 2012) and fit similar models to above. Hazard ratios were then visualised with forest plots created using ggplot2.

## Results

### Development Time

Development was significantly influenced by breeding status, temperature regime and the interaction between the two (χ^2^ (1) = 5.25, *p* = 0.022, χ^*2*^ (1) = 480.75, *p* <0.001 and χ^2^ (1) = 6.49, *p* = 0.011, respectively; Fig. 1). In particular, outbred individuals had a significantly quicker development time in comparison to inbred individuals (Outbred = 21.5 days; Inbred = 22.1 days; *Difference* = −0.56, *p* <0.001, Fig. 1, Table S1A). As expected, individuals at elevated temperatures developed quicker than those in the control regime (Control = 23.0 days; Elevated = 20.6 days; *Difference* = −2.44, *p* <0.001, Fig. 1, Table S1A). Importantly, the detrimental effects of increased mutation load were exacerbated at higher temperatures (OC (22.8 days) – IC (23.2 days): *difference* = −0.348, *p =* 0.02; OE (20.2 days) – IE (21.1 days) *difference* = −0.831, *p* <0.001; Fig 1, Table S1B/C).

**Fig. 1.**
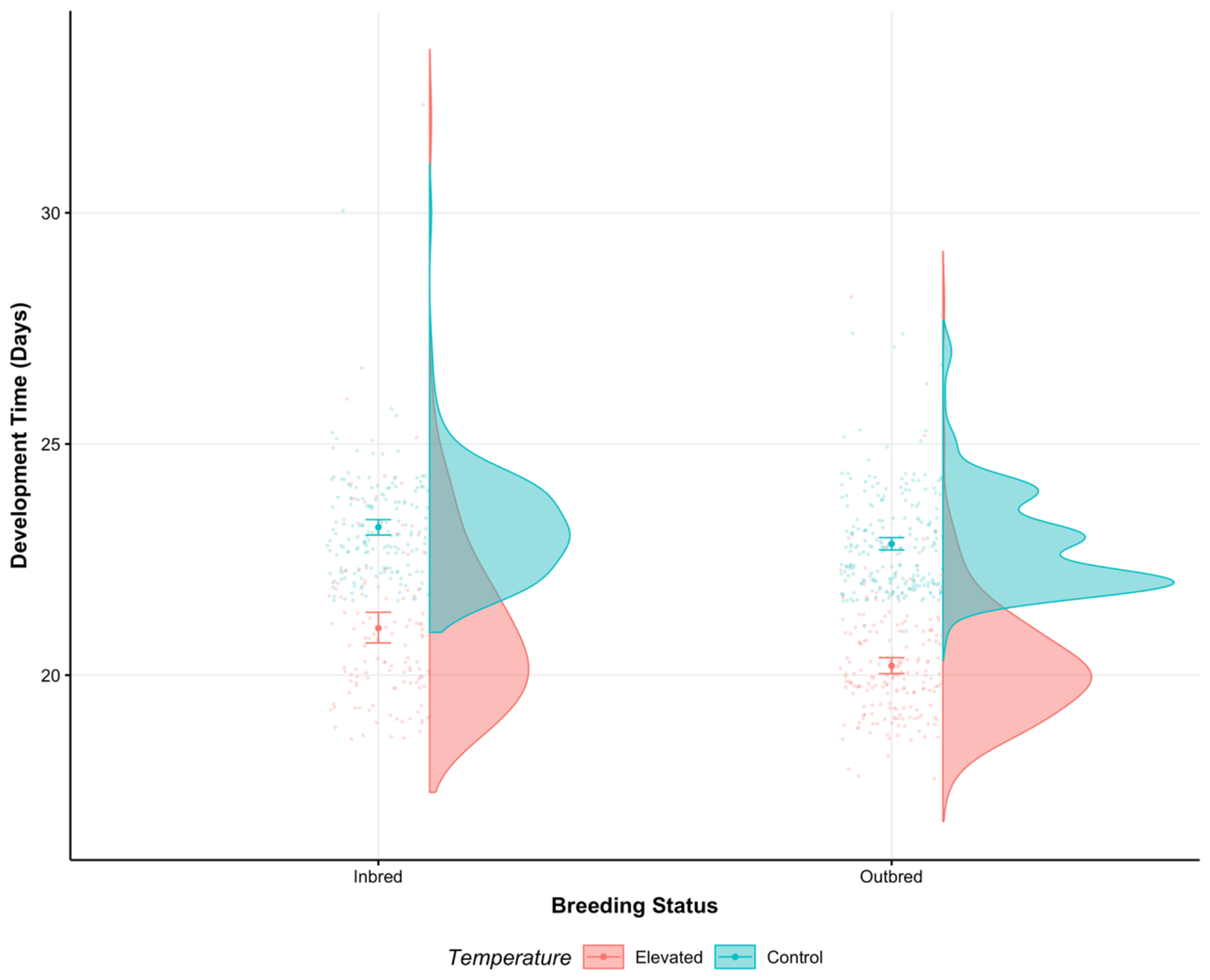
Development time of inbred and outbred populations at control (blue) and elevated (red) temperatures. Points with error bars represent mean values with 95% confidence intervals. Marginal violin plots show the relative distribution of raw data.

### Hatching Success

Hatching success was significantly influenced by breeding status, temperature regime and the interaction between the two (χ^2^ (1) = 35.94, *p* <0.001, χ^*2*^ = 25.09, *p* <0.001, χ^2^ (1) = 3.89 and *p* = 0.049, respectively; Fig. 2). More specifically, hatching success was higher in individuals that were outbred and had a decreased mutation load (Outbred = 87%; Inbred = 59%; *Odds ratio* = 4.50, *p* <0.001; Table S2A) or were exposed to control temperatures and to a less stressful environment (Control = 84%; Elevated = 66%; *Odds ratio* = 2.67, *p* <0.001; Table S2A). This interaction resulted in outbred individuals raised at control temperatures having the greatest hatching success in comparison to other treatments (OC (94%) – OE (78%): *odds ratio* = 4.19, *p* <0.001; IE (50%): *odds ratio* = 14.64, *p* <0.001; IC (68%): *odds ratio* = 6.91, *p* <0.001; Fig. 2; Table S2B/C).

**Fig. 2.**
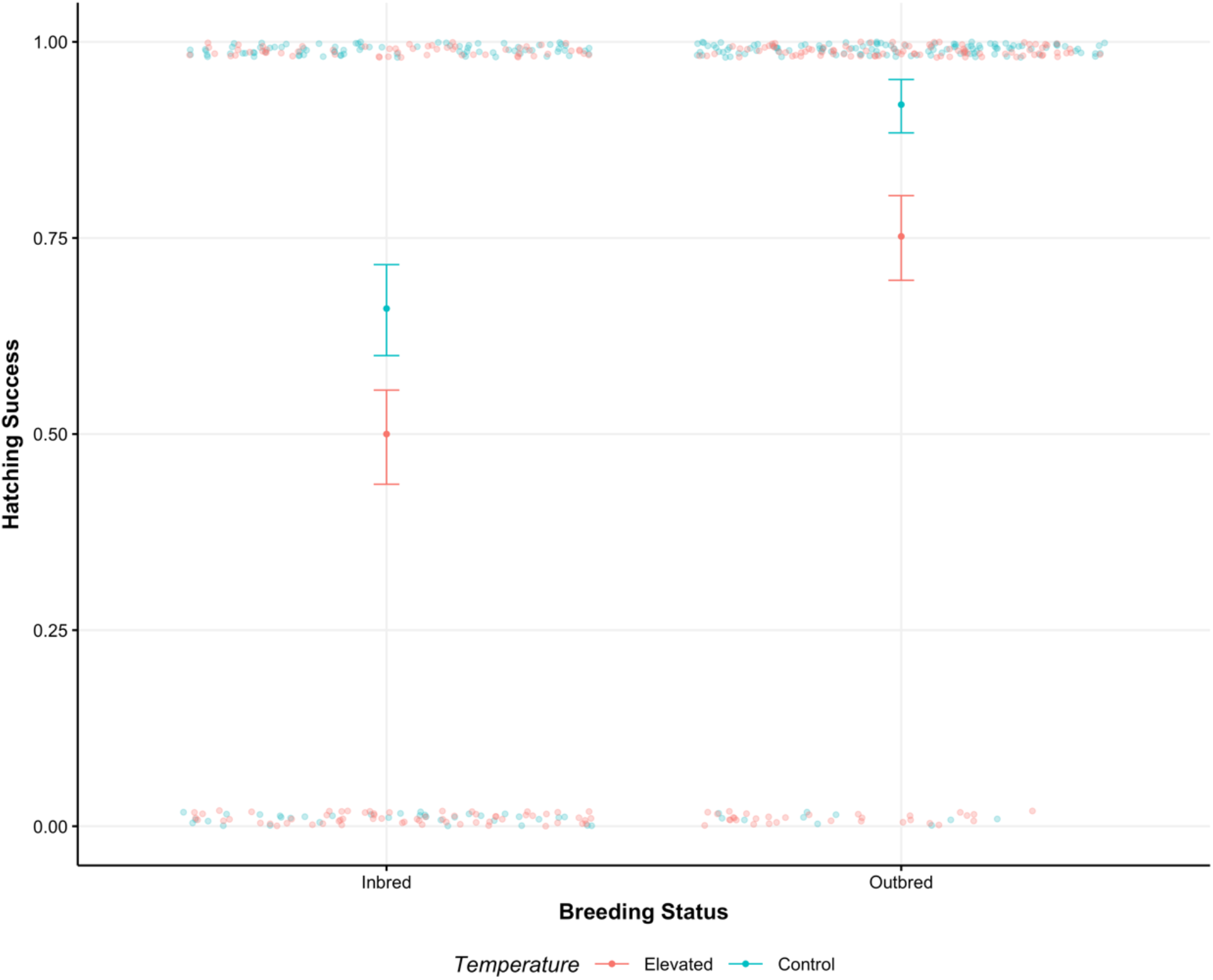
Hatching success in inbred and outbred populations at control (blue) and elevated (red) temperatures. Points between 0 and 1 represent mean values with 95% confidence intervals.

### Reproduction

Age-specific reproduction, LRS and λ*ind* were all significantly influenced by temperature (ASR: χ^*2*^(1) = 5.16, *p =* 0.023; LRS: χ^*2*^ (1) = 279.75, *p* <0.001; λ*ind*: χ^*2*^ (1) = 36.57, *p* <0.001; Fig 3–5, Table S3A-5C) but not breeding status, with no significant difference detected between outbred and inbred individuals (ASR: χ^2^ (1) = 0.003, *p =* 0.958, LRS: χ^*2*^(1) = 0.145, *p =* 0.703, Table; λ*ind:* χ^*2*^ (1) = 0.988, p = 0.320; Fig 3–5, Table S3A-5C). In all cases, elevated temperature was associated with decreased fitness (LRS: Control = 80.0; Elevated = 36.1, *Ratio* = 2.22, *p* <0.001; λ*ind*: Control = 1.13; Elevated = 0.88, *estimate* = 0.25, *p* <0.001; Fig 4–5, Table S4A-5C). Importantly, differences in LRS between inbred and outbred individuals only manifested at elevated temperatures (LRS interaction: χ^*2*^ (1) = 4.08, *p =* 0.043; Fig 4, Table 4A-C), where the negative effects of higher temperatures were exacerbated by increased mutation load (OC (80.7) – IC (79.6): *ratio* = 1.01, *p* = 0.704; OE (38.5) – IE (32.7): *ratio* = 1.18, *p* = 0.02; Fig 4, Table 4A-C). No significant interaction between temperature and breeding status was detected for ASR (χ^*2*^ (1) = 0.429, *p* = 0.512) and λ*ind* (χ^*2*^ (1) = 0.373, *p =* 0.541; Fig 3/5, Table S3A-C/S5A-C).

**Fig. 3.**
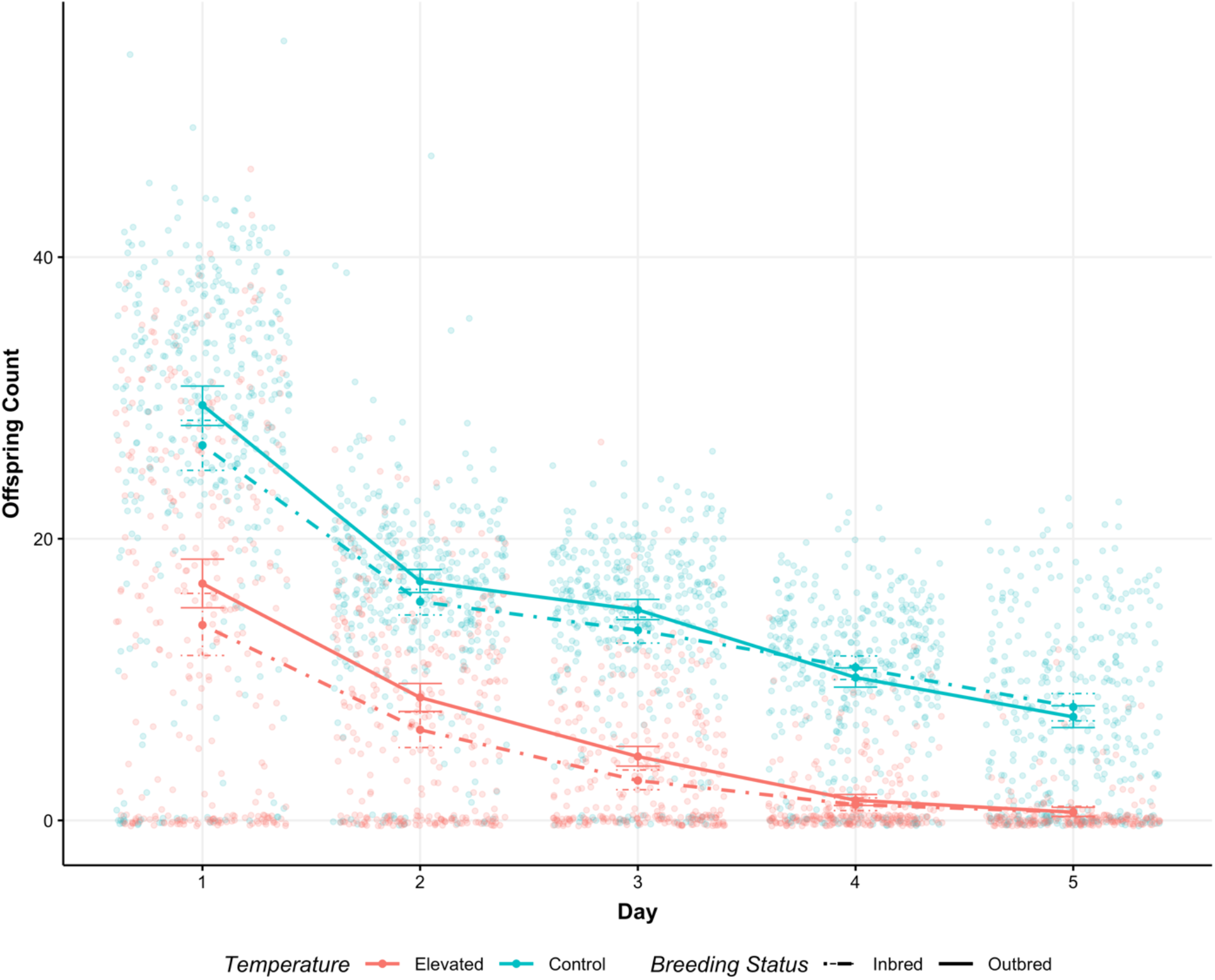
Age-specific reproduction of inbred (solid) or outbred (dot-dash) individuals in elevated (red) or control (blue) temperatures. Points represent means with accompanying 95% confidence intervals.

**Fig. 4.**
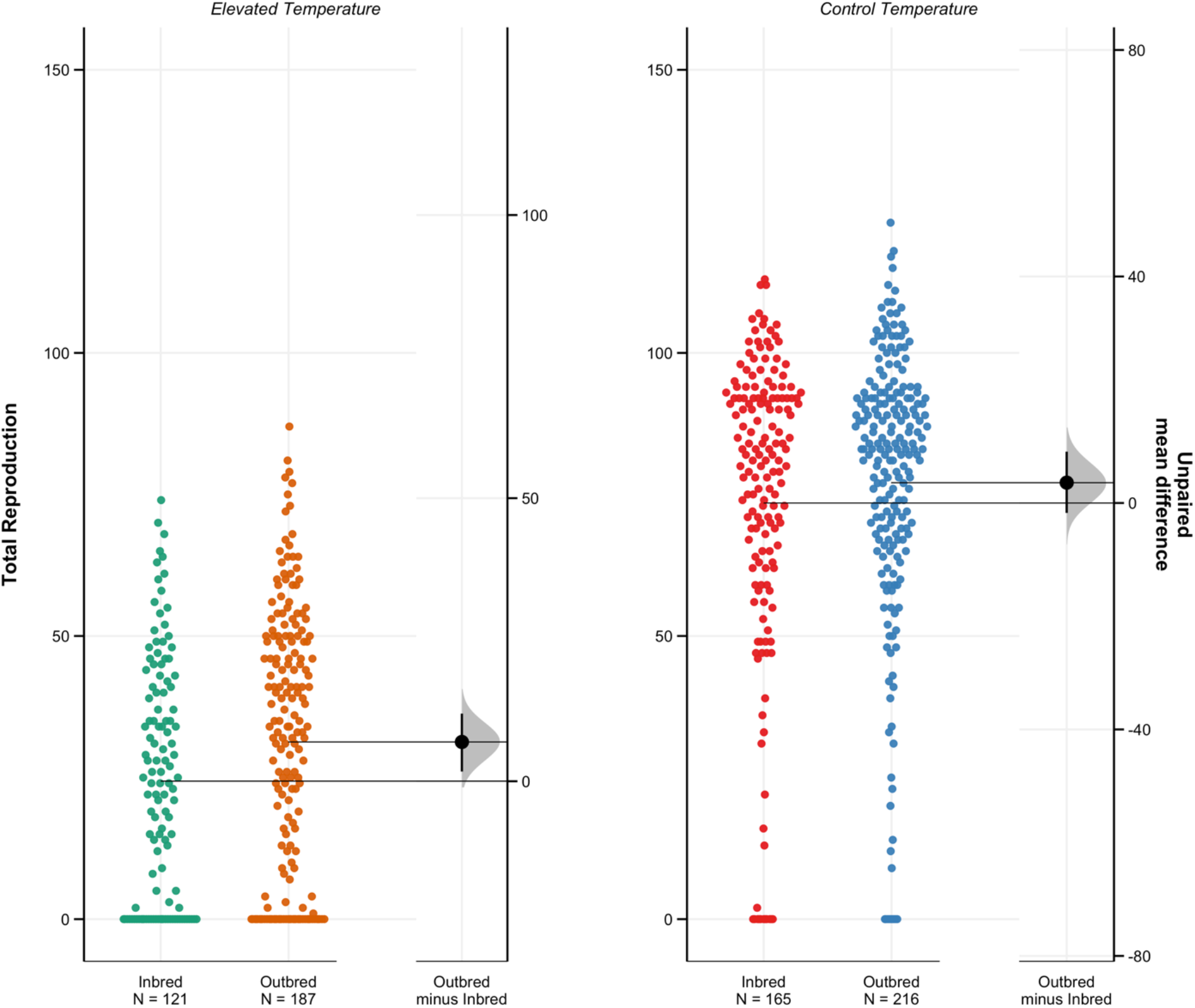
Total reproduction between inbred and outbred individuals at elevated (left) and control (right) temperatures. Each panel shows the raw data and bootstrapped mean differences between treatments with 95% confidence intervals.

**Fig. 5.**
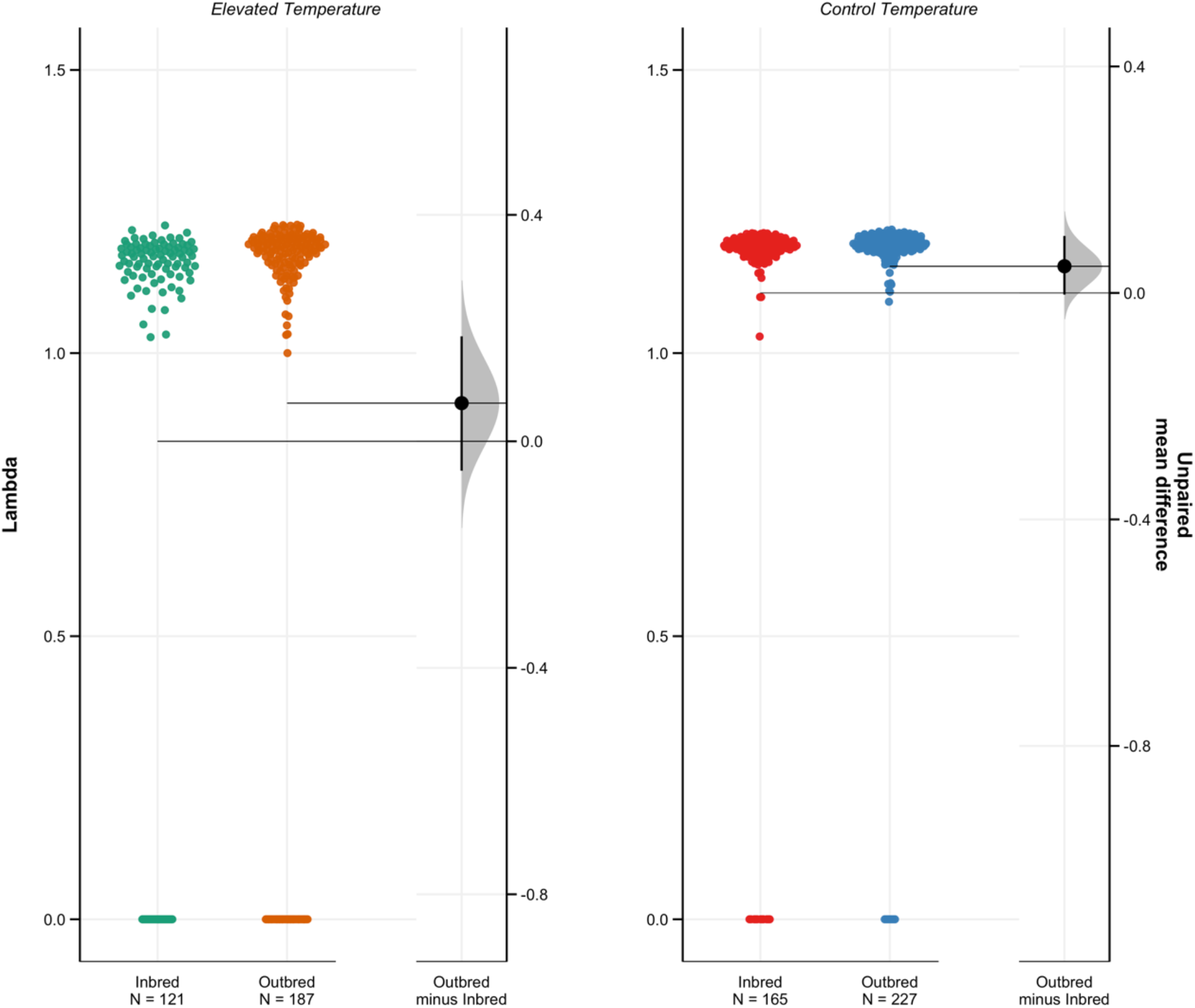
Individual fitness of inbred and outbred individuals at elevated (left) and control (right) temperatures. Each panel shows the raw data and bootstrapped mean differences between treatments with 95% confidence intervals.

### Survival and Longevity

Individual survival and lifespan were significantly affected by temperature regime χ^*2*^ (1) = 183.03, *p*<0.001) but not breeding status (χ^*2*^ (1) = 0.981, *p* = 0.322) or the interaction between the two (χ^*2*^ (1) = 0.929, *p* = 0.335). Individuals raised at control temperatures have reduced mortality risk and longer lifespans in comparison to those from elevated temperatures regardless of breeding status (OC-IC: *estimate* = 0.012, *p* = 0.906; OC-OE: *estimate* = −1.197, *p* <0.001; OC-IE: *estimate* = −1.030, *p* <0.001; OE-IE: *estimate* = 0.167, *p* = 0.169; Fig. 6/7, Table S6A/B).

**Fig. 6.**
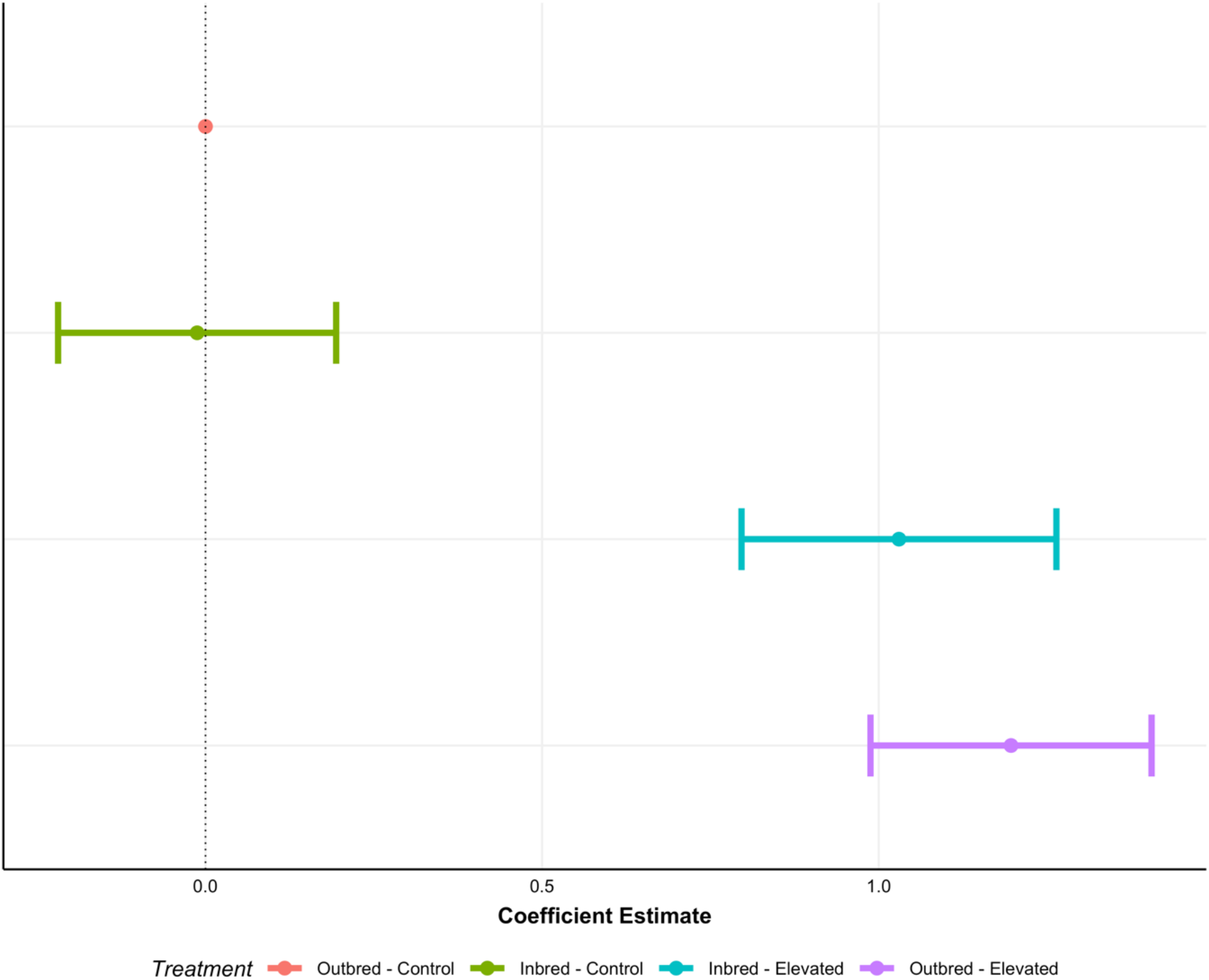
Survival coefficients from a mixed effects cox model with accompanying 95% confidence intervals. Outbred-Control is the reference value at 0. Values to the left reflect mortality decrease and increased longevity, values to the right represent a mortality increase and decreased longevity.

**Fig. 7.**
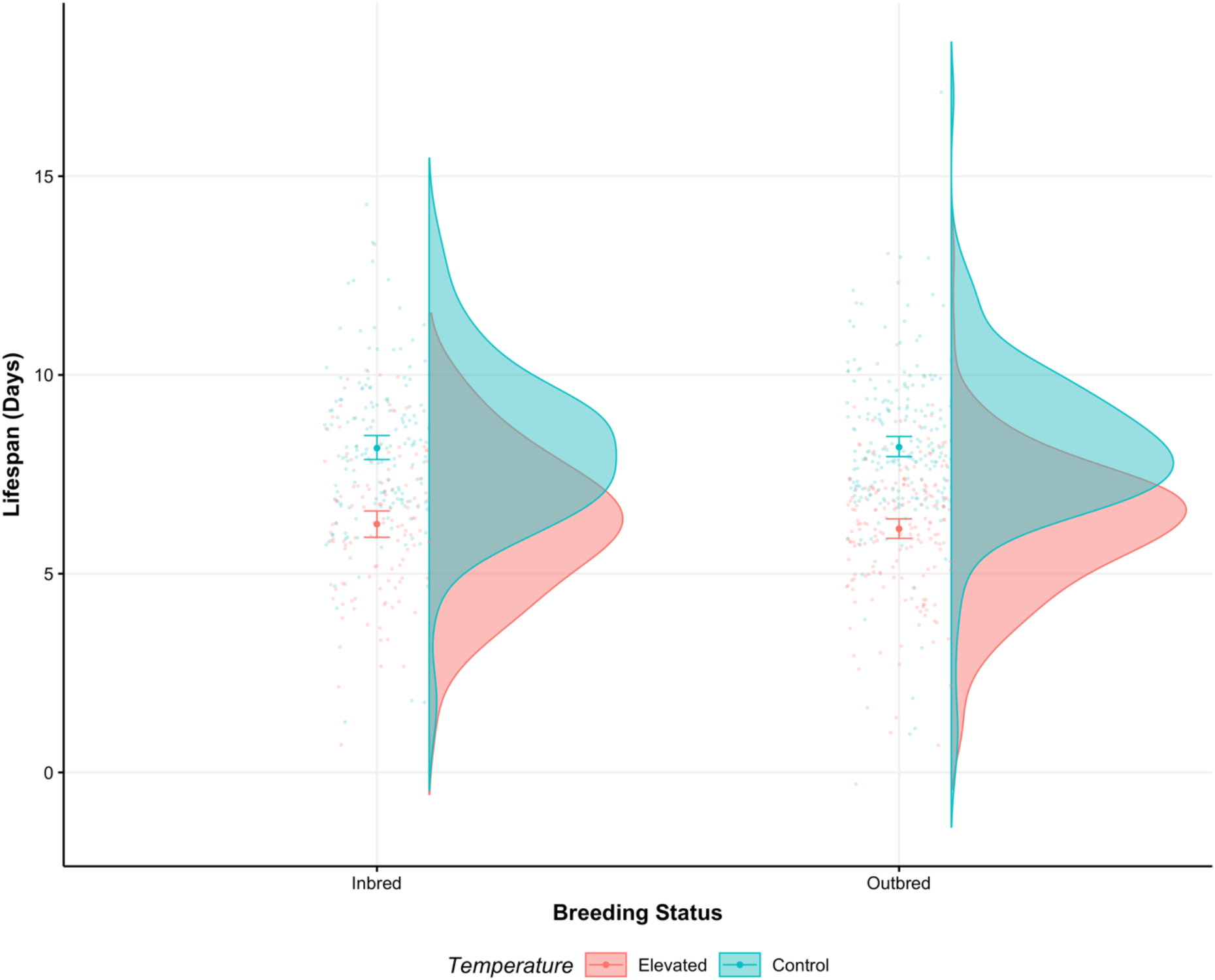
Lifespan of inbred and outbred populations at control (blue) and elevated (red) temperatures. Points with error bars represent mean values with 95% confidence intervals. Marginal violin plots show the relative distribution of raw data.

## Discussion

Our results provide compelling evidence to suggest that increased mutation load from inbreeding exacerbates the negative effects of elevated temperature on various measures and components of fitness. Specifically, we found that increasing temperature and thus exposure to environmental stress had large negative effects on five out of the six measured life history traits. This result alone is unsurprising, as previous work in the same species of beetle has reported similar detrimental effects of high temperature, including reduced reproductive fitness and longevity (Rogell *et al.*, 2014; Berger *et al.*, 2017), but also on the increase of genome-wide *de novo* mutations (Berger *et al.*, 2020). Only on development time was the effect of increasing temperature less obvious. On one hand, faster development time with elevated temperature could be seen as adaptive, as earlier breeding positively influences rate-sensitive fitness (Sibly & Calow, 1986). Under some circumstances, this could compensate for reduced LRS by increasing λ*ind*. However, as this measure (λ*ind*) is entirely dependent on the amount of pre-reproductive time prior to fertility, the variation in development time due to temperature (ranging from 18-32 days) has little impact on the individual fitness value calculated (see Green & Painter, 1975). This is perhaps one reason why we do not see as great a difference in λ*ind* as with LRS. On the other hand, faster development could also be maladaptive, particularly if the increased growth rate results in higher mortality and reduced body size, which will contribute to reduced fecundity (Sibly *et al.*, 1985; Sibly & Calow, 1986). Only when a species becomes adapted to a particular thermal regime (Rogell *et al.*, 2014) or if the environment between parents and offspring is predictable (Sibly & Calow, 1986; Lind *et al.*, 2020) do the harmful effects of increasing temperature begin to subside. However, in contrast to previous work (Yun & Agrawal, 2014), we found that thermal stress, in the absence of any form of density dependence, was sufficient in magnitude to result in increased inbreeding depression on development time, hatching probability and, lifetime reproductive success.

This result is similar in trend to the positive correlation between developmental stress and larval mortality found in several studies in the same organism (Fox & Reed, 2011; Springer *et al.*, 2020). Similarly, a recent study by Springer *et al.* (2020) found a negative effect on female mass, a proxy for female fecundity; however, this was only present within an interaction with another variable, beetle host plant. Nevertheless, in this study we show that *lifelong* stress (i.e. not simply confined to the developmental period) can significantly and detrimentally influence fitness through a reduction in lifetime reproductive success in addition to increasing larval mortality. In addition, we also present another form of inbreeding-environment interaction, in which the control temperature of 29°C also produced significantly reduced hatching probability in inbred individuals which mirrors results from previous work (Fox *et al.*, 2011).

Why such inbreeding-environment interactions should produce deleterious effects on fitness requires an explanation. In a series of elegant studies, Kristensen *et al.* (2002, 2005, 2006) found that inbred *Drosophila* flies were disproportionately expressing genes relating to metabolism and stress response in comparison to outbred individuals (Kristensen *et al.*, 2005). In particular, they found that the heat-shock protein (*Hsp70*) was expressed at higher levels in benign laboratory conditions when individuals were inbred. Importantly, the expression of *Hsp70* is associated with severe and detrimental costs to fitness (Krebs & Feder, 1997; Kristensen *et al.*, 2002). Additionally, when inbred flies were exposed to environmental stress through increasing temperature, they again found differential expression of several important metabolic genes in a synergistic fashion (Kristensen *et al.*, 2006). Taken together, these results, coupled with previous research, suggests that the inbred lines here, which were exposed to both genetic and environmental stress, could be expressing a wide variety of genes that ultimately are contributing to reduced fitness. Future research should focus on understanding whether the same candidate loci found in *Drosophila* are expressed in *C. maculatus* when exposed to both environmental and genetic stress.

The exacerbated negative effects we show here, despite the exposure to the reduced environmental stress of the laboratory (Hedrick & Kalinowski, 2000), highlights the severe and detrimental impact that global climatic changes coupled with habitat fragmentation could have on the survivability of small populations. It is therefore critically important for future conservation research to study these inbreeding-stress interactions in more complex environments using natural populations and with a wide variety of stressors.

## Acknowledgements

Financial support for this project was provided by ERC GermlineAgeingSoma 724909 to AAM and by The American University of Paris to ECB. The authors declare no conflicts of interest.

## Data Accessibility Statement

Data will be deposited in Dryad.

