## Supplementary Material for "Inbreeding reduces fitness of seed beetles under thermal stress"

**Table S1A.** Full model output from the mixed effect model for development time without the interaction

| ***Predictors*** | ***Estimates*** | ***std. Error*** | ***Statistic*** | ***p*** |
|---|---|---|---|---|
| (Intercept) | 22.85 | 0.10 | 227.68 | **<0.001** |
| Breeding [Inbred] | 0.35 | 0.15 | 2.29 | **0.022** |
| Temperature [Elevated] | -2.63 | 0.12 | -21.93 | **<0.001** |
| Breeding [Inbred] * Temperature [Elevated] | 0.48 | 0.19 | 2.55 | **0.011** |
| ***Random Effects*** | | | | |
| $\sigma^2$ | 1.48 | | | |
| $\tau_{00\ Parent.ID}$ | 0.18 | | | |
| $N_{Parent.ID}$ | 100 | | | |
| Observations | 708 | | | |

**Table S1B.** Full model output from the mixed effect model for development time with the interaction

| ***Predictors*** | ***Estimates*** | ***std. Error*** | ***Statistic*** | ***p*** |
|---|---|---|---|---|
| (Intercept) | 22.76 | 0.09 | 242.10 | **<0.001** |
| Breeding [Inbred] | 0.56 | 0.13 | 4.40 | **<0.001** |
| Temperature [Elevated] | -2.44 | 0.09 | -26.09 | **<0.001** |
| ***Random Effects*** | | | | |
| $\sigma^2$ | 1.50 | | | |
| $\tau_{00\ Parent.ID}$ | 0.17 | | | |
| $N_{Parent.ID}$ | 100 | | | |
| Observations | 708 | | | |

**Table S1C.** Pairwise comparisons of relevant treatment contrasts using emmeans

| ***Contrast*** | ***Estimate*** | ***std. Error*** | ***df*** | ***p*** |
|---|---|---|---|---|
| *OC - IC* | -0.348 | 0.152 | 702 | 0.0223 |
| *OE - IE* | -0.831 | 0.166 | 702 | <0.001 |

**Table S2A.** Full model output from the mixed effect model for hatching success without the interaction

| ***Predictors*** | ***Log-Odds*** | ***std. Error*** | ***Statistic*** | ***p*** |
|---|---|---|---|---|
| (Intercept) | 2.38 | 0.21 | 11.45 | **<0.001** |
| Breeding [Inbred] | -1.50 | 0.23 | -6.60 | **<0.001** |
| Temperature [Elevated] | -0.98 | 0.16 | -6.11 | **<0.001** |
| ***Random Effects*** | | | | |
| $\sigma^2$ | 3.29 | | | |
| $\tau_{00\ Parent.ID}$ | 0.58 | | | |
| $N_{Parent.ID}$ | 100 | | | |
| Observations | 1000 | | | |

**Table S2B.** Full model output from the mixed effect model for hatching success with the interaction

| ***Predictors*** | ***Log-Odds*** | ***std. Error*** | ***Statistic*** | ***p*** |
|---|---|---|---|---|
| (Intercept) | 2.68 | 0.27 | 9.84 | **<0.001** |
| Breeding [Inbred] | -1.93 | 0.32 | -6.00 | **<0.001** |
| Temperature [Elevated] | -1.43 | 0.29 | -5.01 | **<0.001** |

| | | | | |
|---|---|---|---|---|
| Breeding [Inbred] * Temperature [Elevated] | 0.68 | 0.35 | 1.97 | **0.049** |
| ***Random Effects*** | | | | |
| $\sigma^2$ | 3.29 | | | |
| $\tau_{00\ Parent.ID}$ | 0.58 | | | |
| $N_{Parent.ID}$ | 100 | | | |
| Observations | 1000 | | | |

**Table S2C.** Pairwise comparisons of relevant treatment contrasts using emmeans

| ***Contrast*** | ***Estimate*** | ***std. Error*** | ***df*** | ***p*** |
|---|---|---|---|---|
| *OC - OE* | 4.186 | 1.196 | 995 | <0.001 |
| *OC - IE* | 14.637 | 4.723 | 995 | <0.001 |
| *OC - IC* | 6.912 | 2.229 | 995 | <0.001 |

**Table S3A.** Model Selection for Age-Specific Reproduction. Showing the top five models in order of AIC with accompanying levels of zero-inflation (*zi*) and dispersion (*di*).

**Significant zero-inflation was identified in a Poisson model w/wo an OLRE (Zero-inflation: *p* <0.001*)***

| *Family* | *Zero-inflation formula* | *zi* | *di* | *AIC* | *df* |
|---|---|---|---|---|---|
| **genpois** | **Breeding*Day + Day2 + Breeding*Temperature** | **1.008** | **0.952*** | **18436.74** | **22** |
| genpois | Breeding*Day + Breeding*Day2 + Breeding*Temperature | 1.009 | 0.952 | 18438.37 | 23 |
| genpois | Breeding*Temperature + Day*Temperature + Day2 | 1.008 | 0.947 | 18442.14 | 22 |
| genpois | Breeding*Day + Day2 + Day*Temperature | 1.007 | 0.949 | 18442.57 | 22 |
| genpois | Breeding*Day2 + Day*Temperature | 1.008 | 0.949 | 18442.71 | 22 |

**Table S3B.** Full model output from the best identified model for Age-Specific Reproduction

| ***Predictors*** | ***Log-Mean*** | ***std. Error*** | ***Statistic*** | ***p*** |
|---|---|---|---|---|
| (Intercept) | 3.82 | 0.05 | 77.41 | **<0.001** |
| Temperature [Elevated] | -0.24 | 0.11 | -2.27 | **0.023** |
| Breeding [Inbred] | 0.00 | 0.08 | 0.05 | 0.958 |
| Day | -0.49 | 0.04 | -12.23 | **<0.001** |
| Day2 | 0.03 | 0.01 | 4.58 | **<0.001** |
| Temperature [Elevated] * Breeding [Inbred] | 0.12 | 0.18 | 0.66 | 0.512 |
| Temperature [Elevated] * Day | -0.02 | 0.11 | -0.17 | 0.866 |
| Breeding [Inbred] * Day | -0.06 | 0.06 | -0.89 | 0.374 |
| Temperature [Elevated] * Day2 | -0.05 | 0.02 | -2.21 | **0.027** |
| Breeding [Inbred] * Day2 | 0.02 | 0.01 | 1.53 | 0.127 |
| (Temperature [Elevated] * Breeding [Inbred]) * Day | -0.23 | 0.18 | -1.26 | 0.207 |
| (Temperature [Elevated] * Breeding [Inbred]) * Day2 | 0.02 | 0.04 | 0.42 | 0.677 |
| ***Zero-Inflated Model*** | | | | |
| (Intercept) | -3.76 | 0.31 | -12.14 | **<0.001** |
| Breeding [Inbred] | 1.44 | 0.35 | 4.16 | **<0.001** |
| Day | -0.08 | 0.19 | -0.42 | 0.677 |
| Day2 | 0.11 | 0.03 | 3.44 | **0.001** |
| Temperature [Elevated] | 2.58 | 0.17 | 15.57 | **<0.001** |
| Breeding [Inbred] * Day | -0.26 | 0.08 | -3.09 | **0.002** |
| Breeding [Inbred] * Temperature [Elevated] | -0.75 | 0.24 | -3.09 | **0.002** |
| ***Random Effects*** | | | | |
| $\sigma^2$ | 0.00 | | | |

| | |
|---|---|
| $\tau_{00\ ID:Parent.ID}$ | 0.05 |
| $\tau_{00\ Parent.ID}$ | 0.00 |
| $N_{ID}$ | 699 |
| $N_{Parent.ID}$ | 100 |
| Observations | 3286 |

**Table S4A.** Model Selection for Total Reproduction. Showing the top five models in order of AIC with accompanying levels of zero-inflation (*zi*) and dispersion (*di*).

**Significant zero-inflation was identified in a Poisson model w/wo an OLRE (Zero-inflation: *p* <0.001*)***

| *Family* | *Zero-inflation formula* | *zi* | *di* | *AIC* | *df* |
|---|---|---|---|---|---|
| genpois | Temperature + Breeding | 1.006 | 0.877 | 5946.163 | 9 |
| **genpois** | **Temperature** | **1.006** | **0.876** | **5946.778** | **8** |
| genpois | Temperature * Breeding | 1.007 | 0.877 | 5947.819 | 10 |
| nbinom2 | Temperature + Breeding | 0.992 | 0.826 | 5961.584 | 9 |
| nbinom2 | Temperature | 0.994 | 0.825 | 5962.198 | 8 |

**Table S4B.** Full model output from the best identified model for Total Reproduction without the interaction.

| ***Predictors*** | ***Log-Mean*** | ***std. Error*** | ***Statistic*** | ***p*** |
|---|---|---|---|---|
| (Intercept) | 4.41 | 0.02 | 182.03 | **<0.001** |
| Temperature [Elevated] | -0.80 | 0.04 | -22.38 | **<0.001** |
| Breeding [Inbred] | -0.05 | 0.03 | -1.42 | 0.157 |
| ***Zero-Inflated Model*** | | | | |
| (Intercept) | -2.95 | 0.24 | -12.52 | **<0.001** |
| Temperature [Elevated] | 1.83 | 0.27 | 6.78 | **<0.001** |
| ***Random Effects*** | | | | |
| $\sigma^2$ | 0.20 | | | |
| $\tau_{00\ Parent.ID}$ | 0.01 | | | |
| $N_{Parent.ID}$ | 100 | | | |
| Observations | 689 | | | |

**Table S4C.** Full model output from the best identified model for Total Reproduction with the interaction.

| ***Predictors*** | ***Log-Mean*** | ***std. Error*** | ***Statistic*** | ***p*** |
|---|---|---|---|---|
| (Intercept) | 4.39 | 0.03 | 172.53 | **<0.001** |
| Temperature [Elevated] | -0.74 | 0.04 | -16.73 | **<0.001** |
| Breeding [Inbred] | -0.01 | 0.04 | -0.38 | 0.704 |
| Temperature [Elevated] * Breeding [Inbred] | -0.15 | 0.07 | -2.02 | **0.043** |
| *Zero-Inflated Model* | | | | |
| (Intercept) | -2.95 | 0.24 | -12.52 | **<0.001** |
| Temperature [Elevated] | 1.83 | 0.27 | 6.78 | **<0.001** |
| ***Random Effects*** | | | | |
| $\sigma^2$ | 0.20 | | | |
| $\tau_{00\ Parent.ID}$ | 0.01 | | | |
| $N_{Parent.ID}$ | 100 | | | |
| Observations | 689 | | | |

**Table S4D.** Pairwise comparisons of relevant treatment contrasts using emmeans

| ***Contrast*** | ***Estimate*** | ***std. Error*** | ***df*** | ***p*** |
|---|---|---|---|---|
| *OC - IC* | 1.01 | 0.0386 | 681 | 0.704 |
| *OE - IE* | 1.18 | 0.0786 | 681 | 0.015 |

**Table S5A.** Full model output from the mixed effect model for individual fitness without the interaction

| ***Predictors*** | ***Estimates*** | ***std. Error*** | ***Statistic*** | ***p*** |
|---|---|---|---|---|
| (Intercept) | 1.15 | 0.02 | 48.51 | **<0.001** |
| Breeding [Inbred] | -0.06 | 0.03 | -1.87 | 0.062 |
| Temperature [Elevated] | -0.25 | 0.03 | -8.38 | **<0.001** |
| ***Random Effects*** | | | | |
| $\sigma^2$ | 0.15 | | | |
| $\tau_{00}$ ParentID | 0.00 | | | |
| N ParentID | 100 | | | |
| Observations | 689 | | | |

**Table S5B.** Full model output from the mixed effect model for individual fitness with the interaction

| ***Predictors*** | ***Estimates*** | ***std. Error*** | ***Statistic*** | ***p*** |
|---|---|---|---|---|
| (Intercept) | 1.15 | 0.03 | 43.39 | **<0.001** |
| Breeding [Inbred] | -0.04 | 0.04 | -0.99 | 0.320 |
| Temperature [Elevated] | -0.23 | 0.04 | -6.05 | **<0.001** |
| Breeding [Inbred] * Temperature [Elevated] | -0.04 | 0.06 | -0.61 | 0.541 |
| ***Random Effects*** | | | | |
| $\sigma^2$ | 0.15 | | | |
| $\tau_{00}$ ParentID | 0.00 | | | |
| N ParentID | 100 | | | |
| Observations | 689 | | | |

**Table S6A.** Full model output from the mixed effects cox model

| ***Predictors*** | ***Estimates*** | ***std. Error*** | ***Statistic*** | ***p*** |
|---|---|---|---|---|
| Breeding [Inbred] | -0.01 | 0.11 | -0.12 | 0.906 |
| Temperature [Elevated] | 1.20 | 0.11 | 11.23 | **<0.001** |
| Breeding [Inbred] * Temperature [Elevated] | -0.15 | 0.16 | -0.96 | 0.335 |
| Observations | 684 | | | |

**Table S6B.** Pairwise comparisons of all other relevant treatment contrasts using emmeans.

| ***Contrast*** | ***Estimate*** | ***std. Error*** | ***p*** |
|---|---|---|---|
| ***OC - IC*** | 0.0124 | 0.105 | 0.9061 |
| ***OC - OE*** | -1.1965 | 0.107 | <.0001 |
| ***OC - IE*** | -1.0300 | 0.119 | <.0001 |
| ***OE - IE*** | 0.1665 | 0.121 | 0.1684 |
